# Effects of mRNA degradation and site-specific transcriptional pausing on protein expression noise

**DOI:** 10.1101/210401

**Authors:** Sangjin Kim, Christine Jacobs-Wagner

## Abstract

Genetically identical cells exhibit diverse phenotypes, even when experiencing the same environment. This phenomenon, in part, originates from cell-to-cell variability (noise) in protein expression. While various kinetic schemes of stochastic transcription initiation are known to affect gene expression noise, how post-transcription initiation events contribute to noise at the protein level remains incompletely understood. To address this question, we developed a stochastic simulation-based model of bacterial gene expression that integrates well-known dependencies between transcription initiation, transcription elongation dynamics, mRNA degradation and translation. We identified realistic conditions under which mRNA lifetime and transcriptional pauses modulate the protein expression noise initially introduced by the promoter architecture. For instance, we found that the short lifetime of bacterial mRNAs facilitates the production of protein bursts. Conversely, RNA polymerase (RNAP) pausing at specific sites during transcription elongation can attenuate protein bursts by fluidizing the RNAP traffic to the point of erasing the effect of a bursty promoter. Pause-prone sites, if located close to the promoter, can also affect noise indirectly by reducing both transcription and translation initiation due to RNAP and ribosome congestion. Our findings highlight how the interplay between transcription initiation, transcription elongation, translation and mRNA degradation shapes the distribution in protein numbers. They also have implications for our understanding of gene evolution and suggest combinatorial strategies for modulating phenotypic variability by genetic engineering.

## Introduction

One of the most important tasks cells do is regulate the level of gene expression—the conversion of genetic information written in DNA into a certain amount of proteins. Interestingly, isogenic cells in the same environment produce variable amounts of mRNA and protein (1-3). Variability (noise) in mRNA and protein levels, however, varies among genes. For example, low noise is expected for genes encoding proteins that are needed in all cells, such as housekeeping proteins. Consistent with this idea, experiments in *Escherichia coli* and budding yeast have shown that genes essential for viability tend to exhibit lower noise than nonessential genes (4-7). For “noisy” genes, such as some genes involved in stress response, a large variability in protein expression can lead to beneficial traits for the population by generating different cell phenotypes. Such a phenotypic heterogeneity is known to offer a “bet-hedging” strategy for bacterial survival in fluctuating and stressful environments (8-11). It can also be beneficial for cooperative social adaptations through “division of labor” within the cell population (12).

Multiple sources of protein expression noise exist. Intrinsic noise arises due to the stochastic nature of gene expression processes. Extrinsic noise can be produced by cell-to-cell heterogeneity in global factors, including the concentration of RNA polymerases (RNAPs) and ribosomes, cell size and the cell cycle (13). Previous experimental and theoretical studies have identified transcription initiation (i.e., the loading of RNAP onto the promoter region) as a major source of intrinsic noise (e.g., 13-27). Specifically, if transcription initiation occurs randomly at a certain frequency, a mode known as “nonbursty” initiation, the mRNA number at steady state follows a Poisson distribution, which characteristically shows an mRNA Fano factor (variance/mean) equal to 1. In contrast, if the rate of transcription initiation varies over time, such as in pulsatile transcription from a promoter that cycles between active and inactive states, the mRNA levels become more variable among cells (mRNA Fano factors > 1). This ON/OFF model of transcription, referred to as “bursty” initiation, is supported by the observation of pulsatile transcription events in live *E. coli* cells (28) and by the grouping of RNAPs along the rRNA operon in electron micrographs (29). The mRNA Fano factors measured in *E. coli* span from 1 to ∼10, suggesting that both nonbursty and bursty promoters may operate *in vivo* (5, 25, 26).

Cell phenotypes are generally dictated at the protein, rather than mRNA, level. Noise in protein levels is often quantified by the squared coefficient of variation (CV^2^), which is the squared standard deviation divided by the squared mean of the protein number distribution (17). Most current analytical models that calculate the noise in protein levels assume that the kinetic parameters associated with the promoter architecture are the major contributing factors in protein synthesis fluctuations, and therefore ignore transcription elongation dynamics and known dependencies between transcription, translation and mRNA degradation (18-22). Analytical models that include transcription elongation exist, but they only consider limit cases of low transcription initiation rate (30) or constant RNAP elongation speed, (31, 32) in order to neglect RNAP-RNAP interactions (RNAP congestion) during elongation. To examine RNAP traffic, modelers have turned to stochastic simulation-based models. This approach has shown that RNAP traffic caused by RNAP pauses can create mRNA and protein bursts from nonbursty promoters (33-38), highlighting the importance of considering transcription elongation dynamics when studying protein expression noise. However, including transcription elongation dynamics in stochastic gene expression models requires many variables and increases the complexity of the model. For this reason, previous simulation-based models have examined special conditions, leaving out translation, mRNA degradation, and/or bursty transcription initiation (33-38).

Here, we explore how various scenarios of transcription elongation dynamics and mRNA degradation affect the noise initially set by bursty and nonbursty promoters. We developed an integrative stochastic model of bacterial protein expression that includes transcription initiation, transcription elongation, mRNA degradation and translation as well as established dependencies, such as the coupling between transcription and translation (39-41), co-transcriptional mRNA degradation (42, 43), and the ribosome effect on mRNA degradation (44) (Fig. 1A). Simulations of this model identified new regimes of post-transcription initiation dynamics that modify the protein expression noise initially set by the promoter.

## Methods

### Modeling transcription, translation, and mRNA degradation

All steps described in this section (Fig. 1B) were stochastically simulated using the Gillespie algorithm (45). For stochastic transcription initiation from a bursty promoter, we generated a series of time points when the promoter was ON or OFF, assuming that the ON and OFF periods follow exponential waiting time distributions with average τ_ON_ and τ_OFF_, respectively. In the case of a nonbursty promoter, the promoter was assumed to be always ON. Next, we determined a series of time points for RNAP loading attempts during ON periods, assuming exponentially distributed waiting times between loading attempts (average rate *k*_loading_). Transcription elongation was modeled by stochastic 1-base pair (bp) stepping, based on the totally asymmetric exclusion process (TASEP) algorithm (46-48). The DNA templates were considered as one-dimensional lattices, where each lattice site corresponded to 1 bp. The stepping rate as a function of template position was provided as an input. For pause-free elongation, we used an average speed of *k*_elongation_. When appropriate, a different stepping rate (inverse of the pause duration, *t*_pause_) was assigned at a pause site (*x*_pause_) for all or a fraction of RNAPs (pausing probability, *P*_pause_). We assumed an exponential dwell time distribution at each nucleotide position based on previous experimental observations (49-51). Steric hindrance between RNAPs was checked before each stepping, assuming an RNAP footprint of 35 bp (52). Similar to previous elongation models (33, 34, 53), we did not include RNAP cooperation upon collision (54) because the kinetics of this process remain unknown. We assumed transcription termination at the end of the template to be instantaneous, though, if desired, slower RNAP release can be modeled by using a slower stepping rate at the last position of the template. After RNAP trajectories were simulated, the spacing between adjacent RNAPs was calculated as time “headways” at every nucleotide position along the gene. Headway is defined as the time interval between two consecutive RNAP exit events at every nucleotide position. Mathematically, it is the subtraction of trajectories of two subsequent RNAPs at a given position (*t*_N_(*x*)-*t*_N-1_(*x*), where *t_N_*(*x*) is the trajectory of the N-th RNAP along the gene). The distribution of headways is considered as an important characteristic of traffic flow (55) and has previously been used to analyze RNAP traffic (48).

**Figure 1.**
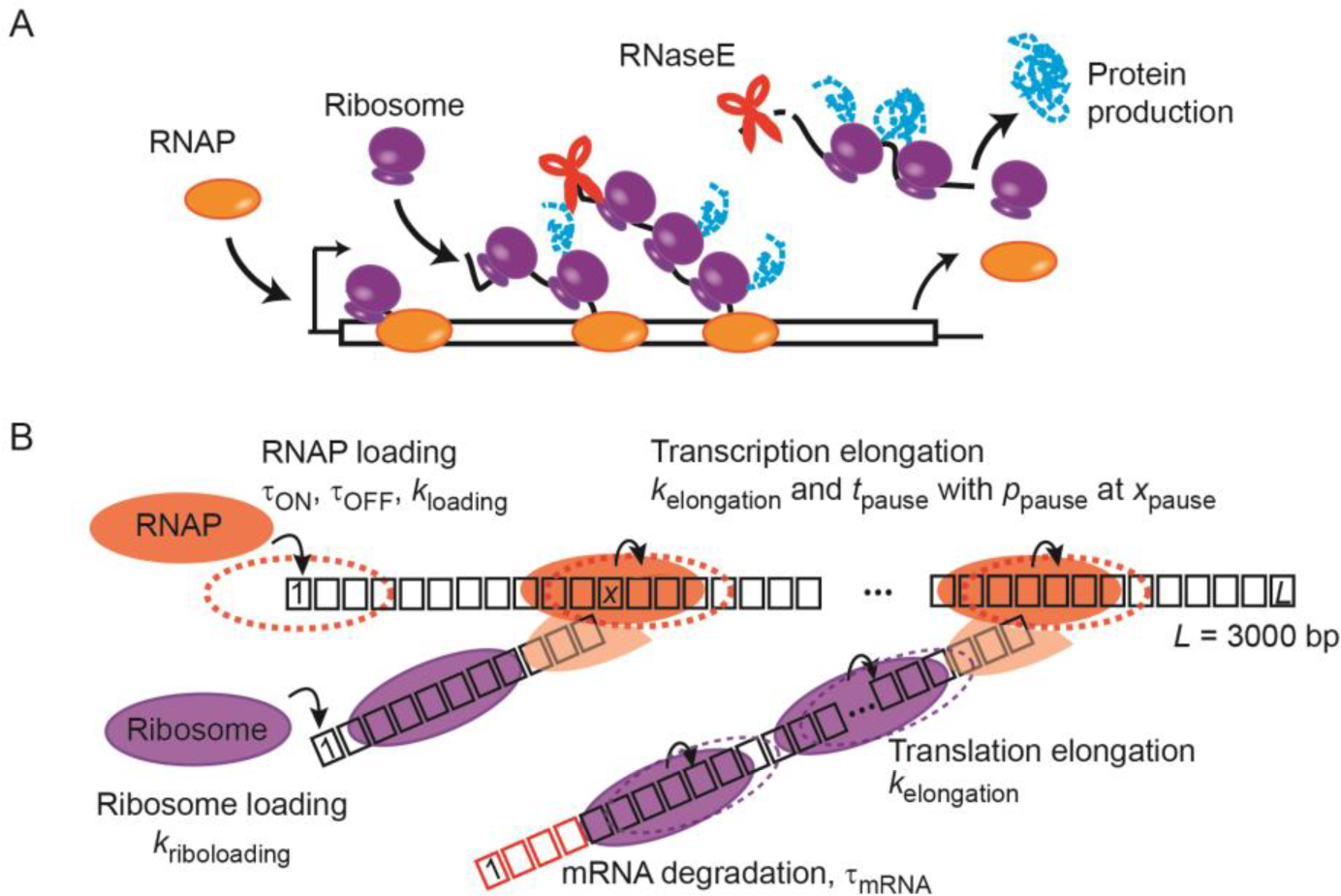
Integrated model of gene expression. (A) Schematic showing the current view on the temporal coordination between transcription, translation, and mRNA degradation in *E. coli*. RNAP loads onto the promoter. Once the ribosome-binding site (RBS) is transcribed, ribosomes load and translocate. mRNA degradation by RNase E can start on the nascent mRNA. The first ribosome maintains contact with the RNAP throughout elongation. In our model, the first full-length protein is produced from an mRNA almost immediately after the RNAP reaches the end of the DNA template. (B) Schematic showing the different steps included in our TASEP-based model of bacterial gene expression. Transcription starts at the first base pair of the template, and transcription elongation occurs by RNAP stepping along the 3,000-bp template. Translation initiates at the first base of the mRNA template (considered here as the RBS), and translation elongation occurs by ribosome stepping on each nucleotide along the mRNA template. mRNA degradation starts from the 5’ end of the mRNA and continues concomitant to the motion of the last ribosome on the transcript. Degraded ribonucleotides are shown in red. Input parameters of our model are indicated.

To model the coupling between transcription and translation, the first ribosome was loaded upon transcription of the first 33 nucleotides (nt). This accounts for the footprint sizes of the RNAP and the ribosome, 35 bp and 30 nt, respectively (52, 56). The first ribosome then moved on the nascent mRNA in concert with the RNAP to maintain contact throughout transcription elongation (39-41). Additional ribosomes were stochastically loaded based on an exponential waiting time distribution (average rate *K*_riboloading_). These ribosomes made stochastic 1-nt steps at the average speed (*K*_elongation_) to reflect the experimental evidence that the average speed of RNAP and ribosomes match (40, 57). During ribosome translocation, the same steric hindrance principle as for RNAP translocation was used: ribosomes were unable to bypass each other on an mRNA, and the first ribosome was unable to bypass the transcribing RNAP on the nascent mRNA.

In our model, mRNA degradation began at the 5’ end of each mRNA, assuming an exponential waiting time distribution between initial synthesis and degradation (with an average lifetime, *τ*_mRNA_). Once the first nucleotide degraded, further ribosome loading was prevented, and the remaining nucleotides on the mRNA were removed concomitant with the movement of the last ribosome on the transcript (58). This model is consistent with experimental observations of 5’-to-3’ net directionality of mRNA degradation (42, 59, 60) and for the ribosome shielding effect (44). In most simulations, protein degradation was considered negligible because most bacterial proteins are very stable (61). However, in some cases, protein degradation was added to the model, assuming first-order kinetics and an average protein lifetime of *τ*_protein_, as previously done (e.g., 36). The whole gene expression process (transcription, translation and mRNA degradation) was simulated for a total duration of 40 min. mRNA levels reached steady state within 10 min of the simulation time under the parameters we used. We performed a total of 1000 simulations for each scenario.

### Analysis of the simulated data

We counted 1 mRNA when the first nucleotide (5’ end) was present, unless noted otherwise. To obtain steady-state distributions of mRNA numbers, we counted the number of mRNAs made but not yet degraded at *t* = 30 min of simulation time (an arbitrary choice of time when mRNA levels were in steady state). We used this distribution to calculate the mean and Fano factor values for each mRNA distribution.

Protein number increased by 1 every time a ribosome reached the end of a transcript. To obtain distributions of protein numbers, we summed all proteins made from a DNA template between *t* = 20-30 min of simulation time, which ensures steady state in mRNA levels. This method of counting proteins is equivalent to measuring protein accumulation over a period of time. Means and CV^2^ of protein numbers were calculated from these distributions.

All error bars indicate an estimation of the standard error of the mean calculated by bootstrap resampling of the original sample size (1,000 simulations) 3,000 times.

## Results

### Comparison between nonbursty and bursty promoters with similar effective transcription initiation rates

While our integrated model of gene expression (see Methods) can be applied to any gene, we modeled the expression of the 3075-nt *lacZ* gene of *E. coli*, which is a popular model in quantitative gene expression studies. Given an average RNAP speed (*K*_elongation_) of 30 nt/sec on the *lacZ* region (Fig. S1A in the Supporting Material) (40), the input average RNAP dwell time at each base position *x* was 1 nt/*K*_elongation_ = 1/30 sec. We used the experimentally determined mean *lacZ* mRNA lifetime of 90 sec (Fig. S1B) as the first-order rate constant for 5’-end degradation (*τ*_mRNA_). For transcription initiation, we varied the RNAP loading rate on the DNA template to achieve a range of expression levels seen in experiments (25). For translation initiation, we used an experimentally-derived average rate of ribosome loading (*K*_riboloading_) of 0.2 sec^-1^ (62, 63).

To build on previously known promoter properties, we considered two different types of promoters: nonbursty and bursty. While the *lac* promoter is thought to be bursty (25, 64), we also considered nonbursty conditions for comparison and to expand our approach to other promoters. The complex, multi-step kinetics of transcription initiation (65, 66) was approximated as one rate-limiting step, as previously done (e.g., 25, 26). Transcription initiation from nonbursty promoters was modeled as a Poisson process with an RNAP loading rate, *K*_loading_, which is the inverse of the loading interval, τ_loading_ (Fig. 2A). This parameter was varied to obtain an output RNAP loading interval between 3 and 500 sec, reflecting the decreasing strength of a constitutive promoter. For the bursty case, the promoters cycled between ON and OFF states, with rate constants *k*_ON_ (rate of switching from OFF to ON state, 1/τ_OFF_) and *k*_OFF_ (rate of switching from ON to OFF state, 1/τ_ON_). RNAPs loaded only during the ON state at an interval of Tloading (RNAP loading interval during the ON state) (Fig. 2A) (22-25). This model resulted in multiple RNAP loading events clustered in time in a pulsatile manner. We used experimentally derived τ_OFF_ = 143 sec and τ_loading_ = 2.2 sec (25), and varied the fraction of time spent in the ON state (*f*_ON_ = τ_ON/(τ__ON_ + τ_OFF_)) from 0.005 to 0.99 to achieve an output average RNAP loading intervals between 3 and 500 sec.

When we compared nonbursty and bursty promoters of similar strength (i.e., yielding similar effective transcription initiation rates and numbers of mRNAs at steady state), we found expected differences at intermediate RNAP loading intervals (e.g., 7, 15 and 30 sec in Fig. 2B-D). First, bursty promoters showed pronounced bursts of RNAP loading events followed by notable OFF periods, resulting in temporal profiles of RNAP trajectories that were very different from those obtained from a nonbursty promoter of similar strength (Fig. 2B). Second, the distribution of RNAPs on the DNA templates was wider for bursty promoters than for nonbursty promoters, with a noticeable peak close to zero due to the stochastic occurrence of OFF and ON states (Fig. S2A). Third, the steady-state distributions of mRNA numbers for bursty initiations were broader, despite having similar mean mRNA numbers (Fig. 2C and Fig. S2B). Fourth, the distribution of “headways”, which is defined by the time interval between two adjacent RNAPs passing a given DNA position (55), appeared largely exponential for nonbursty promoters (Fig. S2C). In contrast, RNAPs from bursty promoters displayed either small headways arising from loading events within an ON period, or large headways arising from loading events separated by an OFF period (Fig. S2C).

**Figure 2.**
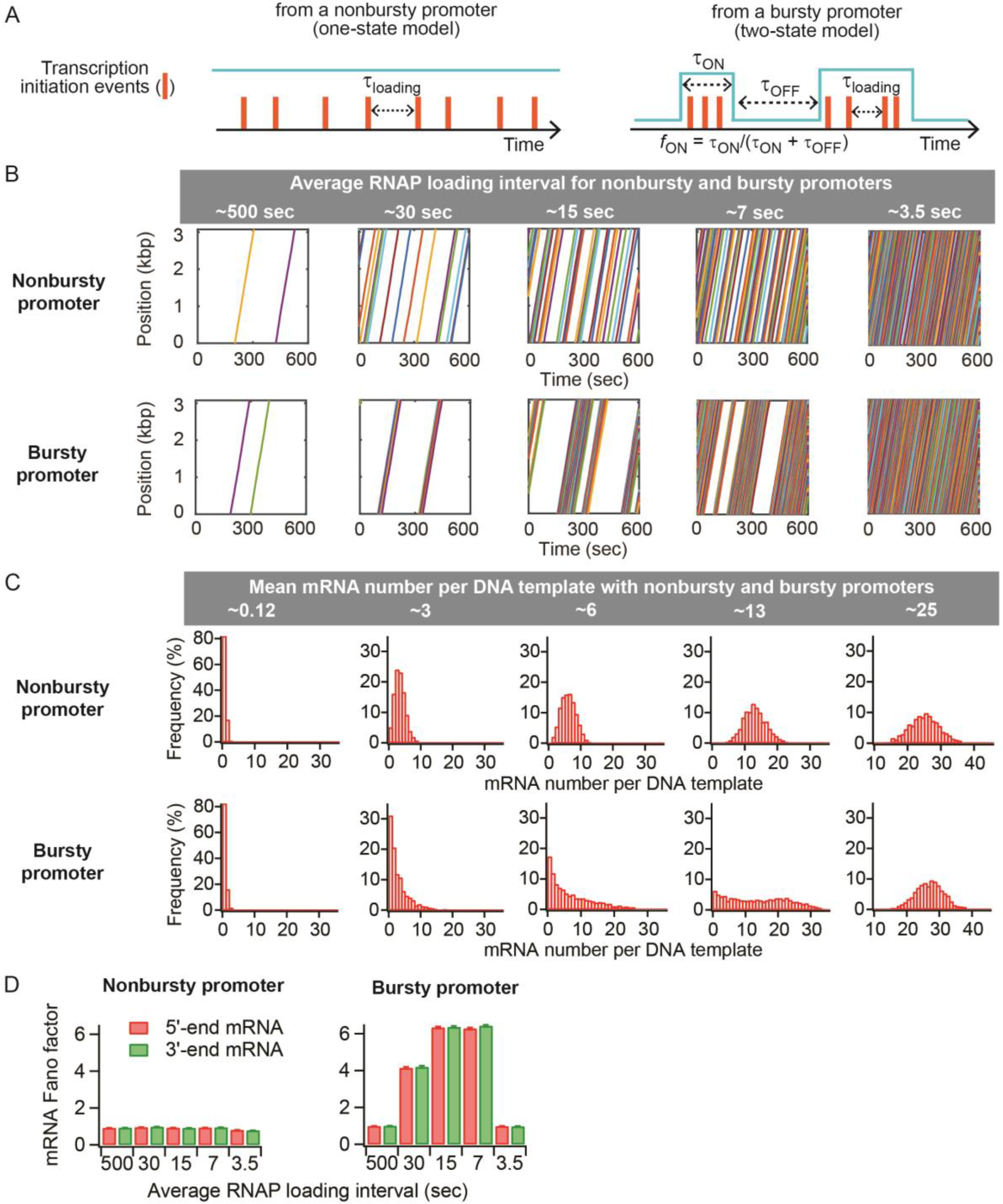
Effect of nonbursty and bursty promoters on RNAP traffic and mRNA number distribution. (A) Schematics for two modes of transcription initiation. The promoter states are shown in cyan, and RNAP loading events are represented by orange bars. (B) Example trajectories of RNAP loading and translocating on individual DNA templates over 10 min based on different transcription initiation conditions: for nonbursty transcription initiation, the input τ_loading_ was varied as 500, 30, 15, 6, 2.5 sec (from left to right) and for bursty transcription initiation, *f*_ON_ was varied as 0.005, 0.1, 0.25, 0.5, 0.99 (from left to right). The average loading interval (grey box) was calculated from the simulation model output. (C) Steady-state distributions of mRNA numbers per DNA template under the transcription initiation conditions used in (B). mRNAs were counted based on the presence of the first base. See Fig. S2B for the results using an mRNA counting method that better reflects how mRNAs are often quantified in fluorescence *in situ* hybridization experiments. (D) Fano factors calculated from the 5’ and 3’ends ofthe mRNAs under the various transcription initiation conditions used in (B-C).

In both promoter cases, the headway set at transcription initiation was conserved until the end of transcription, as shown by the near-perfect overlap in distributions between headways at initiation and termination (Fig. S2C). The conservation of the promoter-dependent “burstiness” until the end of transcription elongation was also shown by comparing the Fano factors calculated from the 5’ and 3’ ends of the mRNA. For both promoter types, the 3’-end mRNA Fano factor remained the same as the 5’-end mRNA Fano factor (1 for nonbursty promoters and greater than 1 for bursty promoters) (Fig. 2D). While a previous modeling study (34) suggested that RNAP bursts set by a bursty promoter can completely disappear during transcription elongation (i.e., 3’-end mRNA Fano factor = 1), we found that such a phenomenon appears when (i) RNAPs are loaded back-to-back during the ON period, and (ii) the RNAP footprint size is modeled as 1 bp (as in ref. 34), instead of 35 bp (Fig. S3). With the smaller RNAP size, many RNAPs load back-to-back during a given ON period, augmenting RNAP congestion and headway separation due to extensive interactions (Fig. S4). We expect that, under realistic parameter values, the memory of a bursty promoter's ON/OFF switch is largely maintained throughout transcription elongation, at least in the absence of RNAP pauses.

At transcription initiation frequencies that were either very low or very high (e.g., average loading interval ≈ 500 sec or 3.5 sec), bursty promoters were virtually indistinguishable from nonbursty promoters (Fig. 2B-D and Fig. S2). At very low initiation frequencies, the ON period was too short to accommodate enough RNAP loading events to exhibit transcriptional bursts. This is consistent with the experimentally determined mRNA Fano factor of 1 for the repressed *lacZ* promoter (25). At very high initiation frequencies, the OFF period was so short that it became negligible (3). These results suggest that transcriptional bursts are unlikely for genes at either side of the expression spectrum.

Importantly, our simulations identified a dynamic range of transcription initiation rates for which our model produced a clear difference between nonbursty and bursty promoters (Fig. 2 and Fig. S2). When we examined protein production under this range of transcription initiation rates, the temporal profile of protein production was largely dictated by the temporal profile of RNAP loading onto the promoter. Nonbursty transcription initiations yielded a relatively constant number of proteins made from a DNA template over time (Fig. 3A). In contrast, bursty transcription initiations resulted in bursty protein productions, showing periods of time without any new protein production from the DNA template (Fig. 3A). As a result, bursty promoters produced the expected broader protein number distribution in comparison to nonbursty promoters, despite having the same mean protein production (Fig. 3B). We also verified that the noise in protein levels increased with increasing RNAP loading intervals (i.e., decreasing transcription initiation rates) from both promoter types (Fig. S5A), consistent with analytical models (18-22).

**Figure 3.**
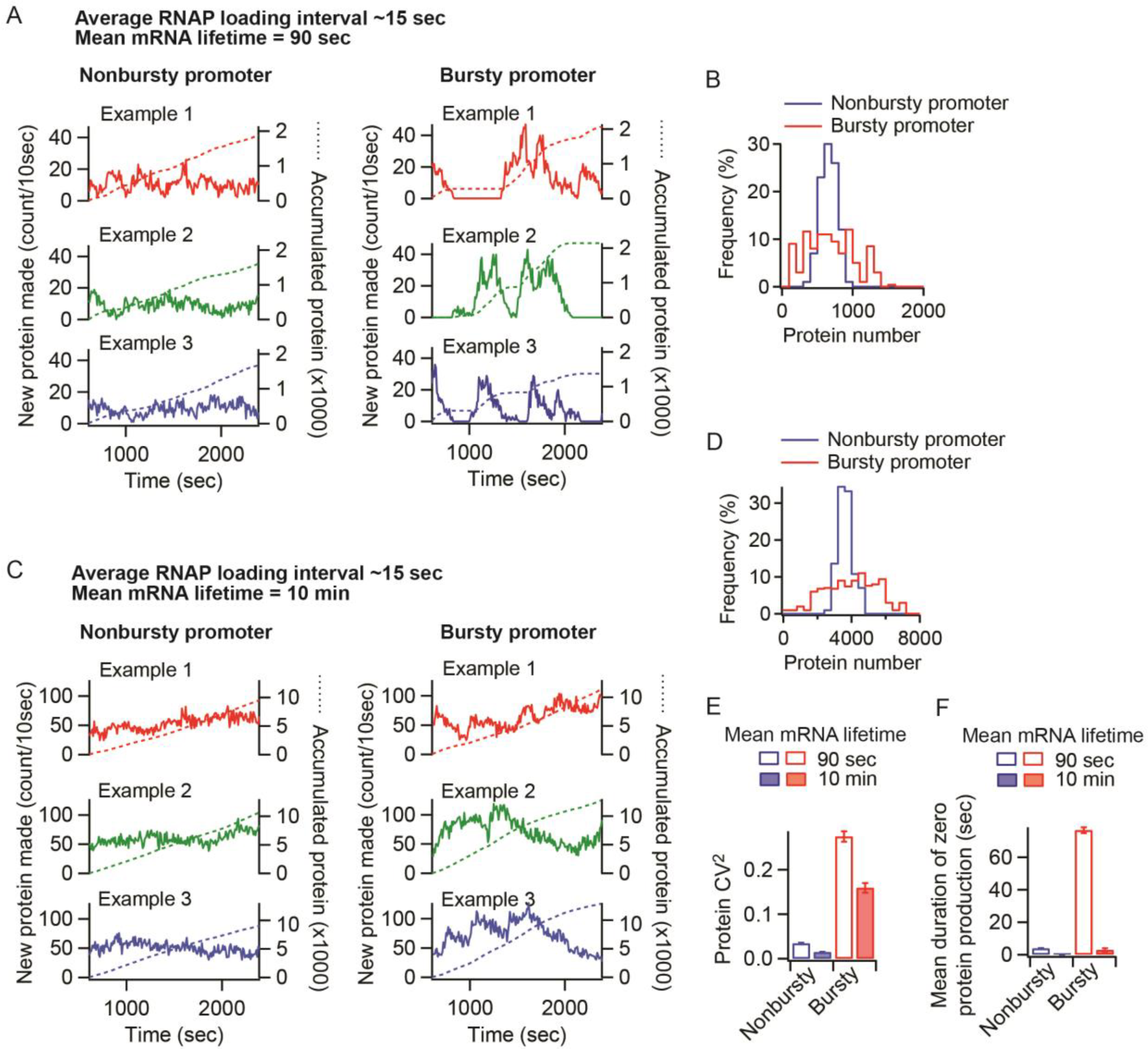
Bursty promoters result in bursty protein synthesis and greater noise in protein levels when the mRNA lifetime is short. (A) Temporal profiles of protein production. The number of new proteins made per DNA template was quantified every 10 sec. Example trajectories were from three independent simulations for nonbursty and bursty promoters with average RNAP loading intervals of ∼15 sec. The mean mRNA lifetime was 90 sec. The number of proteins accumulated from expression of the DNA template is shown in dotted lines. Only the accumulation of proteins occurring within the time frame was calculated. (B) Distributions of the number of proteins made per DNA template over 10 min. (C) Temporal profiles of protein production under the same conditions as in (A), except that the mRNA lifetime was 10 min. (D) Distributions of the number of proteins made per DNA template over 10 min when the mRNA lifetime was 10 min. (E) Variability in protein numbers for a nonbursty and bursty promoter, depending on the mRNA lifetime. (F) Mean durations of zero protein production in the temporal profiles shown in (A) and (C).

### Short mRNA lifetimes facilitate production of protein bursts

Once we had established parameter conditions that clearly distinguish bursty and nonbursty transcription initiations, our goal was to examine how post-transcription initiation processes may affect the burstiness (or lack thereof) set by the promoter. First, we considered mRNA degradation. While the lifetime of the mRNA is known to impact the amount of protein produced (the mean), its effect on the noise in protein levels (CV^2^) is less clear. If both the mRNA lifetime and the transcription initiation rate were changed to maintain the same average protein level, the change in transcription initiation is the dominant factor affecting noise (Fig. S5B), consistent with a previous report (37). But does the CV^2^ vary when the mRNA lifetime changes independently of the transcription initiation rate? When we applied analytical solutions that consider mRNA degradation, we found that they give different answers; some (19, 20) predict a negative effect while others (21, 22) predict no effect at all (Fig. S6A-B).

Simulations of our model showed that the observation of a bursty promoter leading to bursty protein production (Fig. 3A) was dependent on the use of a 90-sec mRNA lifetime. When the lifetime of the mRNA was increased without changing other parameters, protein bursts generated from bursty promoters became less apparent, as illustrated with a 10-min mRNA lifetime (Fig. 3C). While bursty promoters still produced broader protein number distributions than nonbursty promoters (Fig. 3D), the CV^2^ from both types of promoters was reduced by the increase in mRNA lifetime (Fig. 3E). In both cases, the reduction in protein expression noise was correlated with an overall attenuation of temporal fluctuations of protein synthesis, as evidenced by the virtual disappearance of periods of no protein production (Fig. 3F).

We reasoned that the reduced temporal fluctuations of protein synthesis stemmed from the mRNA lifetime being much longer than the RNAP loading interval, resulting in increased protein production between transcription events. Consistent with this idea, the noise in protein expression for both nonbursty and bursty promoters increased either by shortening the mRNA lifetime for a given average RNAP loading interval or by increasing the average RNAP loading interval for a given mRNA lifetime (Fig. 4A-B). In these simulations, protein degradation was neglected because most bacterial proteins are long-lived (61). However, we obtained similar trends when we included protein degradation in our model and simulated an arbitrary average protein lifetime of 20 min (Fig. S6C-D).

**Figure 4.**
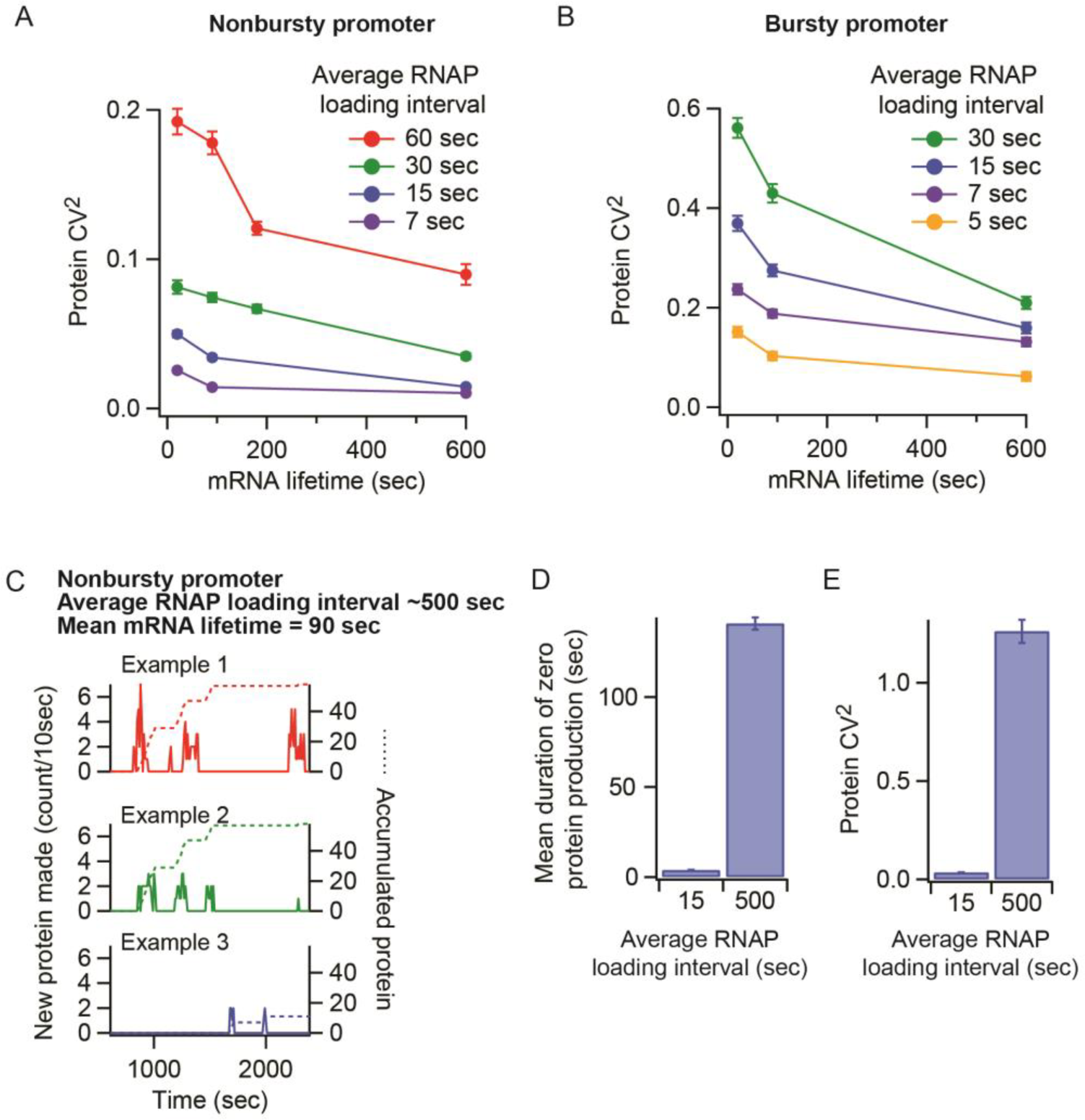
Temporal fluctuations in protein production and variability in protein number are 310 modulated by the RNAP loading interval and the mRNA lifetime. (A and B) Variability in protein numbers as a function of mRNA lifetimes and RNAP loading intervals from nonbursty (A) and bursty (B) promoters. (C) Example temporal profiles of protein production from a nonbursty promoter with an average RNAP loading interval of 500 sec, which is much longer than the 90-sec mRNA lifetime. (D) Mean durations of zero protein production calculated from the temporal profiles of protein production shown in (C) and Fig. 3A. (E) Variability in protein numbers for the conditions described in (D).

When the mRNA lifetime was much smaller than the average RNAP loading interval (e.g., 90 sec vs. 500 sec), a nonbursty promoter was able to produce protein bursts (Fig. 4C and 4D), resulting in higher protein production noise (CV^2^) than when the mRNA lifetime was longer than the average RNAP loading interval (e.g., 90 sec vs. 15 sec) (Fig. 4E). This is consistent with *in vivo* observations that occasional firing of the repressed *lac* promoter (average RNAP loading interval of 40-150 min under the experimental condition used in the cited studies) causes spikes of protein production (62, 67). This is because each mRNA is degraded before the next one is made, resulting in well-separated bursts of protein production.

These results suggest that short mRNA lifetimes (in the minute time scale), a common characteristic of bacterial mRNAs (42, 59, 68), facilitate bursty protein synthesis and increase the variability in protein levels across the population for both bursty and nonbursty promoters.

### Sequence-dependent pauses can attenuate RNAP bursts and reduce protein expression noise

So far, our simulations considered pause-free elongation. In *in vitro* experiments, RNAPs can pause seemingly randomly along the DNA template (49, 69, 70). Modeling studies have shown that long stochastic (sequence-independent) pauses can produce RNAP bursts from nonbursty promoters, because RNAPs can pile up behind the paused RNAP and form a convoy that travels together once the pause ends (33-36). RNAPs are also known to pause at specific DNA sites for durations that generally range from seconds to ∼1 minute (49, 51, 71-80). Pause sites are common in *E. coli* based on RNAP profiling experiments (81, 82). Previous modeling work has shown that sequence-dependent pauses of 100 or 500 sec generate protein bursts from nonbursty promoters due to ribosome piling behind the paused RNAP (37). However, such long-lived RNAP pauses are expected to be rare, and it is unclear whether shorter pauses at specific DNA sites can still affect the noise in protein levels. Furthermore, to our knowledge, the role of sequence-dependent RNAP pauses has only been reported in the case of nonbursty promoters. Whether pause-prone sites affect gene expression noise from bursty promoters is unknown.

In our model of sequence-dependent pausing, RNAPs resided at each nucleotide on average for 1/30 sec, as before, except at the pause site (*x*_pause_), where we varied the average RNAP dwell (*t*_pause_). The probability of a pause at the particular site (*P*_pause_) was also considered, such that if a pause occurred, the dwell time was randomly chosen from an exponential distribution with a mean dwell time of *t*_pause_ (49-51). For illustration purposes, we modeled a single pause site in a 3-kbp gene, driven by either a nonbursty or bursty promoter with an average RNAP loading interval of ∼15 sec. Even at *P*_pause_ = 100%, pauses shorter than the RNAP loading interval, such as *t*_pause_ = 1 sec, had a negligible effect on RNAP traffic regardless of the promoter type, as shown by the near-perfectly overlapping distributions of Δheadways (headway at the end of elongation - headway at the start for two consecutive RNAPs) between the 1-sec pause and the no-pause cases (Fig. S7).

**Figure 5.**
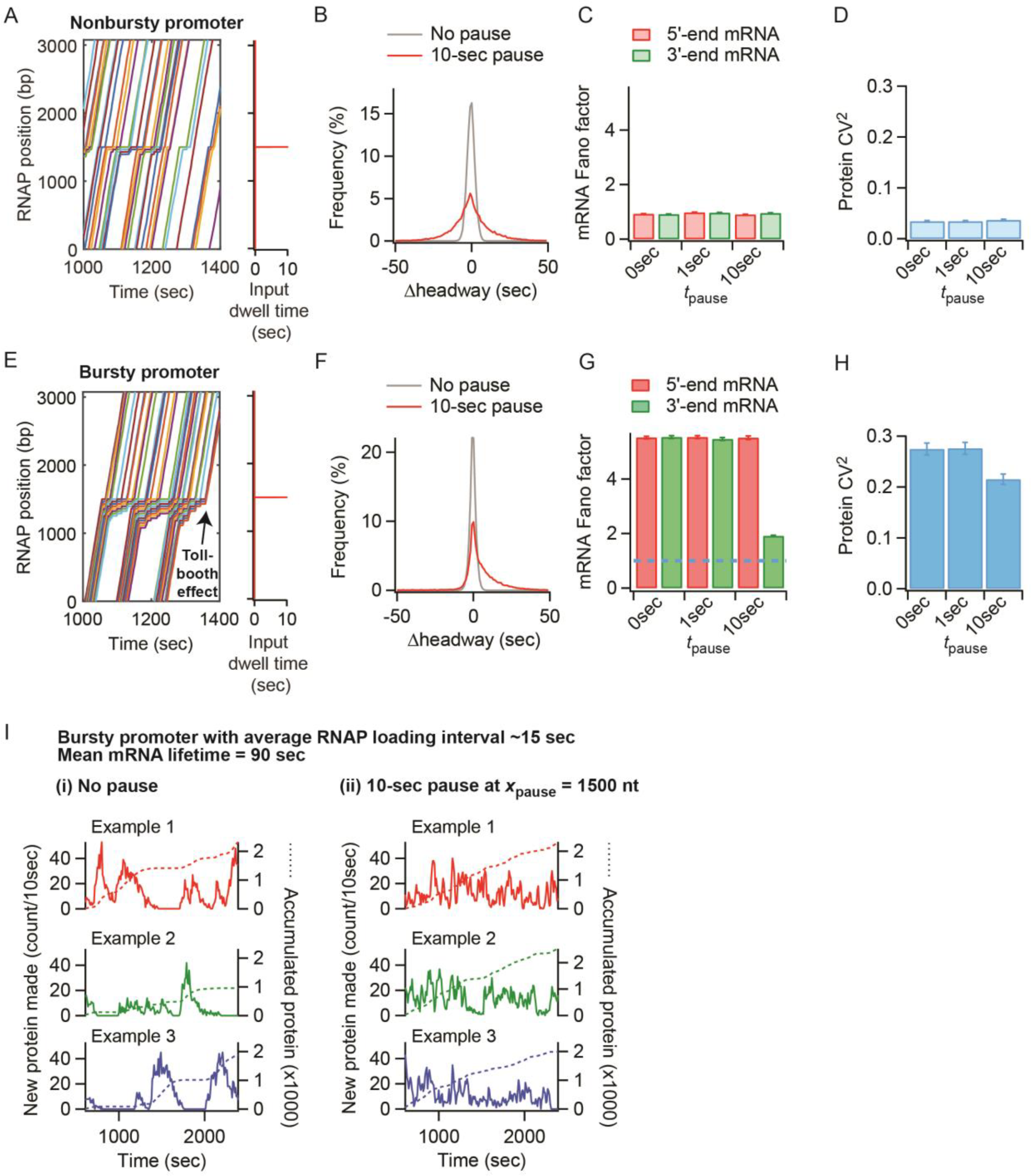
Sequence-dependent RNAP pauses affect gene expression dynamics and steady-state distributions of mRNA and protein numbers. (A) Example trajectories of RNAPs along a DNA template from a simulation with a pause duration tpause = 10 sec at xpause = 1500 nt with ppause = 80% for a nonbursty promoter with average RNAP loading interval of ∼15 sec. (B) Distributions of Δheadways for all simulated RNAPs using the same conditions as in (A). (C) Fano factors calculated based on either the 5’ or 3’ end of the mRNA, considering three different pause conditions for the nonbursty promoter: no pause (0 sec), 1-sec pause at *x*_pause_ = 1500 nt with *P*_pause_ = 100%, and 10-sec pause at *x*_pause_ = 1500 nt with *P*_pause_ = 80%. (D) Protein noise (CV^2^) of the nonbursty promoter, considering three different pause conditions as in (C). (E) Example trajectories of RNAPs along a DNA template with the same pause condition as in (A) but with a bursty promoter and an average RNAP loading interval of ∼15 sec. (F) Distributions of Δheadways from all simulated RNAPs using the same conditions as in (E). (G) 5’- and 3’- end mRNA Fano factors for the bursty promoter with different pause conditions, as used in (C). The blue dotted line denotes a Fano factor equal to 1, representing a nonbursty process. (H) Protein noise (CV^2^) for the bursty promoter, considering three different pause conditions used in (C). (I) Effect of a pause on temporal fluctuations in protein production from a bursty promoter with an average RNAP loading interval of ∼15 sec. While the temporal profiles of protein production in the absence of a pause show bursty protein production (left), those in the presence of a 10-sec pause at *x*_pause_ = 1500 nt with *P*_pause_ = 80% show a reduction in bursty protein production. In particular, the pause smoothens the temporal profile of protein accumulation (dotted lines).

When the pause was similar to (e.g., *t*_pause_ = 10 sec) or longer than the RNAP loading intervals and the probability of pausing was high (e.g., *P*_pause_ ∼ 80%, as for the *his* and *ops* pauses (51, 76, 79)), the pause resulted in two effects on RNAP traffic: (i) RNAP piling upstream of the pause site and (ii) a change in RNAP headway downstream of the pause. In the case of a nonbursty initiation, these effects (Fig. 5A) resulted in a broader distribution of Δheadways between subsequent RNAPs, but the Δheadway distribution remained centered around zero, with no net change (Fig. 5B). The nonbursty conditions remained until the end of the template because the headway between subsequent RNAPs either increased or decreased with similar probabilities. As a consequence, the Fano factor was maintained at ∼1 at both the 5’ and 3’ ends of the mRNA (Fig. 5C), and the noise in protein levels (CV^2^) was not affected (Fig. 5D). Unlike a 100-sec pause (37), a 10-sec pause did not provide sufficient time for ribosomes to pile up behind the paused RNAP to create substantial protein bursts (Fig. S8). In the case of bursty promoters, the frequent back-to-back loading of RNAPs (i.e., small initial headway) caused the majority of RNAPs to catch up with each other at the pause site (Fig. 5E). As most RNAPs in the pile will also stop at the pause site, they will resume traveling with a new headway dictated by the pause duration. As a result, the headway between RNAPs showed a net increase after the pause site (Fig. 5F). This situation is analogous to car traffic near a tollbooth. The congested traffic before the tollbooth becomes fluid after the cars pay the toll. The pause-dependent increase in headway led initial RNAP clusters to largely dissipate into a nonbursty-like situation after the pause, as seen in the example RNAP trajectories (Fig. 5E) and in the large decrease in mRNA Fano factor between 5’ and 3’ ends (Fig. 5G). In other words, pauses similar to or longer than RNAP loading intervals diminished the effect from the bursty promoter's ON/OFF switch by increasing the RNAP headway after the pause. This memory loss of initial conditions lowered protein expression noise (Fig. 5H) by smoothing the temporal profile of protein production (Fig. 5I). Pauses also resulted in a noise-attenuating effect when we considered protein degradation (see Fig. S9 for a protein lifetime of 20 min).

When we examined the effect of imposing two pause sites (*t*_pause_ = 10 sec, *p*_pause_ = 100% and *x*_pause_ = 1,500 and 2,500 bp) on a gene driven by a bursty promoter, we found that the second pause did not affect the RNAP headway as much as the first pause (Fig. S10, scenarios (ii) vs. (i)). We reasoned that the headway increase generated by the first pause reduced the number of RNAPs that pile at the second pause site. However, when the second pause was longer than the first (e.g., *t*_pause_ = 10 and 15 sec, respectively), it further increased RNAP headways (Fig. S10, scenarios (iii) vs. (i)), suggesting that multiple long-lived pauses can have additive effects on RNAP traffic.

## Discussion

In this study, we highlight the role of post-transcription initiation processes, such as mRNA degradation and RNAP pausing, in altering the intrinsic noise in protein expression dictated at transcription initiation by the promoter.

While the lifetime of mRNA is well known to alter the amount of proteins produced, its potential effect on protein noise was less clear based on previous theoretical studies (Fig. S6A-B). We found that mRNA lifetimes longer than the OFF period of a bursty promoter dampen the temporal fluctuations of protein synthesis (Fig. 3A vs. 3C). Because one mRNA typically generates more than one protein, longer mRNA lifetime reduces the effect of the bursty promoter's ON/OFF switch (Fig. 4B). mRNA expression from nonbursty promoters also fluctuates over time due to the stochastic nature of transcription initiation. Hence, the mRNA lifetime also smooths temporal fluctuations of protein production in the case of nonbursty promoters (Fig. 3C and 4A). Altogether, this suggests that mRNA degradation is a factor to consider when studying noise in gene expression, especially given that the lifetimes of bacterial mRNAs can vary over an order of magnitude (42, 59, 68).

RNAP pausing is another important post-transcription initiation event that can affect noise. So far, pauses have been viewed as noise-generating factors (33-38). This work suggests that RNAP pauses can also attenuate noise by modulating RNAP traffic downstream of the pause (Fig. 5E-I). The RNAP headway, which shapes the temporal fluctuations in mRNA and protein production, can be altered by a pause (Fig. 5E-F) to the point that the memory of a bursty promoter's ON/OFF switch can be lost after the pause site (Fig. 5G).

Whether transcriptional pausing attenuates or generates mRNA and protein bursts depends on the probability of RNAPs to stop at a particular DNA site. If an RNAP pause occurs stochastically at a random position along the gene (i.e., low probability of pausing at any given position), it can create a line of RNAPs behind the pause that travels as a convoy when the pause ends (33-36). This is akin to a traffic light situation in which all cars stopped at a red light move together when the light turns green. The size of the RNAP convoy, which dictates the size of mRNA and protein bursts, increases with the duration of the pause, the RNAP loading rate and the RNAP translocation speed. A similar noise-generating effect on RNAP traffic is expected if a low-probability pause occurs at a specific DNA sequence (36, 38). However, if a pause has a high probability of occurring, it has an opposite noise-attenuating effect. Here, as mentioned before, the traffic analogy is with a tollbooth where all cars must stop, one by one, before resuming travel with a new headway. RNAPs accumulated behind the pause have to stop at the pause site before being released one RNAP at a time (Fig. 5E). Emergence from the pause site results in more fluid traffic, diminishing any promoter-induced noise effects (Fig. 5G-H).

*In vitro* single-molecule experiments have shown that the probability of an RNAP pause at a given location correlates with its duration (51). Therefore, sequences that generate pauses long enough to alter RNAP traffic are likely to be efficient at pausing RNAPs, favoring the “tollbooth” noise-attenuating mechanism that we report. That said, low-probability pauses that are long-lived are also likely to occur inside cells. For instance, in *E. coli,* RNAPs occasionally retain the initiation sigma factor, σ^70^, during elongation (83-93), and these σ^70^-associated RNAPs can pause at promoter-like sequences within genes for very long periods of time (minutes) (85, 88, 91, 92, 94). Since a minority of the RNAPs retain σ^70^ after initiation (83-92), only a fraction of the RNAP population will pause, effectively triggering the “traffic-light” noise-generating mechanism described above (Fig. S11). Furthermore, because of the coupling between transcription and translation in bacteria, these long-lived RNAP pauses may also affect the traffic of ribosomes, resulting in sharper protein bursts (Fig. S11).

Location is yet another important pause property to consider. In the *E. coli* genome, RNAP pause sites are often located near promoters (81, 83, 85, 86, 91, 95). RNAP piling behind the pause site may extend to the promoter and prevent the loading of additional RNAPs (53, 95-97). Since the transcription initiation rate inversely scales with CV^2^ (Fig. 4A-B) (15, 16, 19), this promoter blockage can indirectly increase noise (Fig. S12A-B). Furthermore, RNAP stalling near the promoter can also result in reduced translation initiation rates (Fig. S12C) when ribosomes piling on the nascent transcript reach and block the ribosome-binding site (RBS) (Fig. S12D). This second effect on protein expression rate stems from the temporal coupling between transcription and translation in bacteria.

Thus, transcriptional pausing can create opposite effects on noise depending on the pause probability, duration and location. Interestingly, the longevity of a pause site in a *Salmonella* magnesium transporter gene has been shown to change in response to varying concentrations of magnesium (79). This example raises the possibility of environmental regulation of noise by modulating pause duration. Overall, our study stresses the importance of a comprehensive model of gene expression when estimating noise, including for the analysis of genome-wide trends in gene expression noise (3) since promoter architectures and pause properties vary among genes.

As congested traffic dynamics of RNAPs and ribosomes have yet to be solved analytically, a simulation-based model, such as ours, provides a convenient tool for testing different scenarios and for estimating the combinatorial effect of noise modulators. For this purpose, we provide our MATLAB-based simulation code and detailed guidelines in the Supporting Text. Simulations generate mRNA and protein distributions, as well as RNAP and ribosome traffic dynamics. While we tested the model using the _t_ON_, t_OFF_,_ and T_loading_ parameter values reported for the *lacZ* promoter, our model is generalizable and can be extended to genes with different promoter architectures by simply varying τ_ON_, τ_OFF_, and τ_loading_. A limitation of our current model is that it does not include processes, such as physical “pushing” between RNAPs (54) and potential premature termination of transcription and translation at RNAP and ribosome congested sites (98, 99), which may mitigate pause-dependent effects on protein expression noise. Determining the kinetic parameters of these processes will facilitate their integration in future models.

mRNA lifetimes and RNAP pauses are evolvable features at the gene-specific level because they are sequence-dependent and can change through mutations. Our work predicts that mutations altering pause conditions (e.g., duration and probability) or mRNA lifetime (e.g., by altering mRNA secondary structure at the 5’-untranslated region) will affect protein expression noise at the level of individual genes. Our work also suggests possible ways by which protein expression noise may change globally. For example, mutations that render RNAP less prone to pausing (e.g., *rpoB5101* mutation in *E. coli* (76)) are expected to affect the protein expression noise of pause-sensitive genes.

In summary, by comparing bursty and nonbursty transcription initiations under a variety of scenarios, we highlight conditions under which bursty promoters produce nonbursty protein production and nonbursty promoters generate bursty protein profiles. These findings underscore the combinatorial origin of protein expression noise. Noise-modulating factors can have opposite effects depending on parameter conditions. The combinatorial effect of these factors may affect how genome sequences evolve by modulating phenotypic variability within a population. Combinatorial approaches could also be exploited for genetic engineering in synthetic biology. For instance, our findings suggest conditions to maximize protein expression noise and phenotypic diversity: a bursty promoter, a short mRNA lifetime, and an absence of long RNAP pauses. The opposite conditions are expected to minimize phenotypic heterogeneity.

## Supporting Material

Supporting Material contains Supporting Text, Supporting Materials and Methods, twelve figures and one table. The source code can be found in http://github.com/JacobsWagnerLab/TASEPnoise.

## Supporting Citations

References (100-103) appear in the Supporting Material.

## Author Contributions

C.J.-W and S.K. designed the study and wrote the manuscript. S.K. performed the experiments and modeling. S.K. and C.J.-W. analyzed the data. C.J.-W supervised the project.

## Acknowledgements

We thank Bruno Beltran and Drs. Damon Clark, Thierry Emonet and Alvaro Sanchez for helpful discussions. We also thank the members of the Jacobs-Wagner Laboratory for discussions and for critical reading of the manuscript. C.J.-W. is an investigator of the Howard Hughes Medical Institute. The authors declare no conflict of interest.

## Notes

Competing interests: The authors declare that no competing interests exist.

